# GeneCup: mine PubMed for gene relationships using custom ontology and deep learning

**DOI:** 10.1101/2020.09.17.297358

**Authors:** Mustafa Hakan Gunturkun, Efraim Flashner, Tengfei Wang, Megan K. Mulligan, Robert W. Williams, Pjotr Prins, Hao Chen

## Abstract

Interpreting and integrating results from omics studies typically requires a comprehensive and time consuming survey of extant literature. Here, we introduce GeneCup, an easy to use literature mining web service that searches all PubMed abstracts for user-provided gene symbols in conjunction with a set of custom keywords organized into a customized ontology, as well as results from human genome-wide association studies (GWAS). As an example, we organized over 300 keywords related to drug addiction into seven categories. The literature search is conducted by querying the NIH PubMed server using a programming interface, which is followed by retrieving abstracts from a local copy of the PubMed archive. The main results presented to the user are individual sentences containing the gene symbol, organized by the keywords they also contain. These sentences are presented through an interactive graphical interface or as tables. GWAS results are displayed using a similar method. All results are linked to the original abstract in PubMed. In addition, a convolutional neural network is employed to distinguish sentences describing systemic stress from those describing cellular stress. The automated and comprehensive search strategy provided by GeneCup facilitates the integration of new discoveries from omic studies with existing literature. GeneCup is free and open source software. The source code of GeneCup and the link to a running instance is available at https://github.com/hakangunturkun/GeneCup

## 1. Introduction

We describe a web service and application—*Mining <u>gene</u> relationships using <u>cu</u>stom ontology from <u>P</u>ubMed* (GeneCup) (http://genecup.org)—that automatically extracts information from PubMed and NHGRI-EBI GWAS catalog on the relationship of any gene with a list of keywords hierarchically organized into a user created ontology. In addition, genetic associations related to the keywords are also retrieved from the GWAS catalog. As an example, we created an ontology for drug addiction related concepts containing seven categories and over 300 keywords. We will describe the details of GeneCup by using this ontology.

Omic studies are becoming the main driving force for discovering molecular mechanisms of human diseases. Over 5000 genome-wide association studies (GWAS) have mapped over 71,000 associations between genetic variants and diseases/traits (Buniello *et al*. 2019). For example, GWAS has become the main platform of discovery on genetic variants responsible for phenotypes related to substance abuse and psychiatric disorders. One recent human GWAS identified over 500 variants associated with smoking and alcohol usage related traits (Liu *et al*. 2019). GWAS on other drugs of abuse, such as opioid (Polimanti *et al*. 2020) or cocaine (Huggett and Stallings 2020) have also been conducted or are ongoing. GWAS on psychiatric diseases also had numerous successes. A recent survey identified 1223 genome-wide significant SNPs associated with psychiatric phenotypes (Horwitz *et al*. 2019). Many risk SNPs are shared among addiction and psychiatric phenotypes (Horwitz *et al*. 2019). Specialized databases, such as the GWAS catalog (Buniello *et al*. 2019), are available for searching the association between genetic variants and phenotypes. Genetic mapping studies using model organisms, such as worms, flies, mice and rats, have also identified many associations between genetic variants and drug abuse related phenotypes. These phenotypes range from response to or voluntary consumption of cocaine, opioids, nicotine, alcohol, etc. (Engleman *et al*. 2016; Adkins *et al*. 2017; Highfill *et al*. 2019; Zhou *et al*. 2019). Transcriptome (Farris *et al*. 2015b; Lo Iacono *et al*. 2016; Zhang *et al*. 2016; Kapoor *et al*. 2019; Cates *et al*. 2019; Huggett and Stallings 2020) or epigenome (Ponomarev *et al*. 2012; Farris *et al*. 2015a; De Sa Nogueira *et al*. 2019) profiling using bulk tissue or single cells (Avey *et al*. 2018; Karagiannis *et al*. 2020) have also discovered the involvement of many genes in response to drugs of abuse, stress, or other psychiatric related conditions.

In these omics studies, understanding the function of genes is a challenging task that requires thorough integration of existing knowledge. Statistics-driven gene ontology, or pathway analysis, are often employed for this purpose. However, extensive review of the primary literature is ultimately needed to provide a comprehensive and nuanced narrative of these mechanisms. For many scientists, this starts as searches of PubMed based on their domain knowledge. These ad hoc searches often miss important information not only because of the inherent complexity of the biology, but also because of the amount of time required for designing a search strategy, conducting the searches, reading the texts, extracting relevant facts, and organizing them into categories. The task of literature searches is especially daunting when many genes are identified in a single study. To the best of our knowledge, GeneCup is the only web service designed to alleviate much of these manual labors by bringing the relevant facts from PubMed and GWAS catalogue to the users.

GeneCup relies mostly on keyword matching to select relevant sentences. However, as in the example of addiction ontology the same keyword can have multiple meanings. In particular, stress promotes initial drug use, escalates continued drug use, precipitates relapse and is a major factor contributing to drug addiction (Koob and Schulkin 2019). Stress in this context refers to the body’s response to internal and external challenges and is mediated by activating the hypothalamic–pituitary–adrenal axis. In addition, stress can also refer to the responses of cells to perturbations of their environment, such as extreme temperature, mechanical damage, or accumulation of metabolites, etc. These responses often involve the activation of specific molecular pathways. Both systemic and cellular stress have a large collection of literature. We therefore developed a machine learning model to separate sentences describing cellular stress from those that describe system stress.

GeneCup is available as a free web-service. In addition, its source code is available for those interested in setting up a service of their own or modifying the code to better suit their needs.

## 2. Methods

### 2.1. System Overview

GeneCup is a free and open source web application (Fig. 1). The source code and URL of a running instance is available at https://github.com/hakangunturkun/GeneCup. The main user interface contains a search box that accepts up to 200 gene symbols from the user. Each gene symbol is then paired with each one of the custom ontological categories to query PubMed. The title and abstract of these records are then obtained from a mirrored copy of PubMed on the local server. Sentences containing at least one gene symbol and one keyword are retained. A local copy of NHGRI-EBI GWAS catalog is also searched for associations between the queried genes and phenotypes related to the ontology. The results are available as an interactive graph or a table that provides links to key sentences from the abstracts, which in turn, are linked to PubMed. In the example of addiction keywords, sentences that contain the keyword “stress” are further classified into two types (i.e. systemic vs cellular) before presented to the user, by using a one dimensional convolutional neural network.

**Figure 1:**
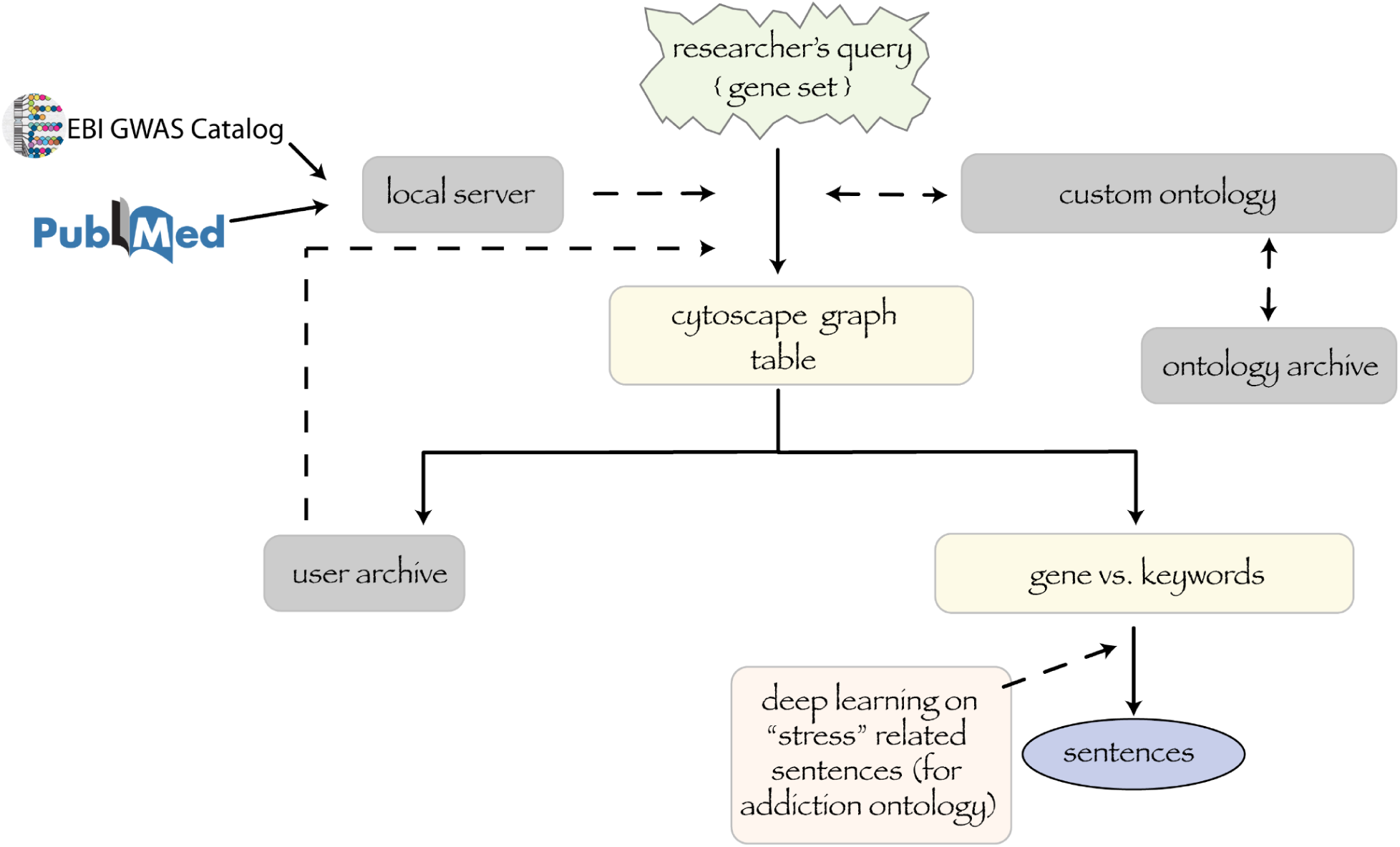
Overview of the workflow of GeneCup. GeneCup allows researchers to query the relationship of any gene with a list of keywords hierarchically organized into a user created ontology. This information is automatically extracted from PubMed and NHGRI-EBI GWAS catalog. The users have an option to choose keyword categories during the search. Searches are conducted using EUtils against the PubMed database but abstracts are retrieved from a locally mirrored copy of PubMed. The results are displayed as a cytoscape graph (Fig. 3) and a table. The graph and the table have many interactive elements, including displaying sentences that include the gene symbols and the keywords. The number of unique abstracts and related sentences are shown separately. Custom ontologies and search results are archived on the server if the user chooses to log in. If addiction ontology is used, sentences containing the keyword *stress* are classified using a convolutional neural network into one of two classes: systemic stress or cellular stress (Figs. 2 and 4).

### 2.2. Sources of data: PubMed and GWAS catalog

We created a copy of the entire PubMed abstract on our server following instructions provided by the NCBI (Kans 2020). This allows us to rapidly retrieve the abstracts and bypass the limits imposed by NCBI on automated retrievals to prevent system overload. This local copy is updated automatically every week on our server.

We also store a local copy of the GWAS catalog database (Buniello *et al*. 2019) (i.e. all associations v1.0.2 from https://www.ebi.ac.uk/gwas/docs/file-downloads). This file is updated manually upon every new release of the catalog. This allows us to perform customized and rapid queries.

### 2.3. User defined ontologies

The custom ontology has three-levels. The top level is the name of the categories, which can be used to decide whether its sub-categories are included in a new search. The second level are concepts that are displayed in the results (i.e. interactive graphs and tables). The third level contains the actual keywords used in PubMed queries and finding matching sentences. For example, the top level “cells” can contain second level concepts such as “neurons” and “glial cells”, with “glial cells” further containing keywords such as “astrocytes”, “microglia”, etc. The matching keywords at the third level are highlighted using bold font when the sentences are displayed. A special top level keyword “GWAS” is reserved for searching the GWAS catalog. Any keyword under this branch is used to search the GWAS catalog database. This is a flexible structure that allows the user to freely organize a large collection of keywords to fit their needs. A free user account is needed for creating and editing custom ontologies.

### 2.4. Query processing and user interfaces

We wrote the web-service in the Python programming language and used the Flask library as the web application framework (“Flask”). Users of the web service have the option of creating an account for the purpose of saving search results for later reviews. Query terms provided by the user are first paired with all the keywords. Keywords belonging to the same second level ontology terms are combined using the boolean OR operator before joining with the gene symbol using the AND operator. The E-utilities provided by the NCBI Entrez system (Kans 2020) are used to send the query to the PubMed database (Esearch) and to retrieve PMIDs (Efetch). Corresponding records for each PMID are obtained from the local copy of PubMed and the xtract tool is used to parse the titles and abstracts. The Python NLTK library (Bird *et al*. 2009) is then used to tokenize the abstracts into sentences. Python regular expressions are used to find sentences that contain at least one instance of a query gene and one instance of a keyword. The number of *abstracts* containing such sentences are then counted. The gene is also searched in the GWAS catalog for phenotypic associations. The number of associations are also counted. A network graph is constructed using the Cytoscape Javascript library (Shannon *et al*. 2003), where all genes, keywords, and GWAS terms are used as nodes, and a connection is made between nodes describing a gene and a keyword. The number of abstracts are used as the weight of the edge. This interactive graph allows a user to click on the edge to review the corresponding sentences. All sentences are linked to their original PubMed abstract. The user can also click on a gene to see its synonyms. These synonyms are obtained from the NCBI gene database but are not included in the original search. This is because they often do not appear in the literature or have other meanings and thus provide inaccurate results. However, the web interface allows these synonyms to be included in a new search to retrieve additional information that is potentially relevant.

We also provide a set of scripts for querying large numbers of genes at the Linux command line. The first script counts the number of relevant abstracts. The second script retrieves the abstracts and extracts the relevant sentences. A third script then generates an html page containing all the results. The intermediate results can be examined and cutoff thresholds can be determined between the scripts. This set of scripts can be executed without a web browser and thus is suitable for querying much larger set of genes (e.g. >1000)

Lastly, queries can also be initiated by placing the terms in the URL. For example, to start a search for CHRNA5 and BDNF genes against the keyword categories drug, stress, addiction, and GWAS, the following hyperlink can be used: https://genecup.org/progress?type=drug&type=stress&type=GWAS&type=addiction&query=CHRNA5+BDNF

This allows links to GeneCup queries to be embedded into other websites. When the hyperlink above is clicked, the results in graphical format will appear in a separate window.

The GeneCup source code is distributed as free and open source software and can therefore easily be installed on other systems. The whole service with dependencies is described as a byte reproducible GNU Guix software package (Wurmus *et al*. 2018).

### 2.3. Mini-ontology for addiction related concepts

We created a mini-ontology for addiction related concepts (Table S1). The top level has the following seven categories: addiction stage, drugs, brain region, CNS cell type, stress, psychiatric diseases and molecular function. The second level is composed of relevant keywords and the third level includes subconcepts of the keywords or commonly used spelling or acronyms for the keywords. Users have the option to skip any category to speed up the query.

### 2.5. Finding the most researched genes related to addiction

Using the script interface described above, we first retrieved all (61,636) human genes from the NCBI gene website (NCBI). We parsed the gene symbols together with their aliases and counted the total number of abstracts for each gene using E-Utils. Relevant sentences for all genes with more than 50 abstracts were then retrieved. We manually examined the most studied 100 genes with the most abstracts iteratively and removed 988 words from the list of gene symbols and aliases before the majority of the sentences in the final results are relevant for the search. Because of the need to manually inspect the results and exclude gene synonyms that yield false matchings, this function is not provided in the web interface.

### 2.6. Convolutional neural network to classify sentences describing stress

There are many machine learning methods that have been applied to natural language processing (NLP) tasks. Young et al (Young *et al*. 2017) compared some of the deep learning related algorithms employed in different types of NLP tasks. Among them, convolutional neural network (CNN) has been shown to be efficient in many sentence level classification projects (dos Santos *et al*. 2015; Francis-Landau *et al*. 2016; Lopez and Kalita 2017; Gehring *et al*. 2017; Wang and Gang 2018). CNN was initially designed for two dimensional image processing (Lecun *et al*. 1998). It uses a linear operation called convolution besides the regular neural network components, and explores the important patterns in a data by identifying both local and global features of the data. The ability to detect nonlinear relationships among the features effectively is one of the key advantages of deep learning architectures. Here, we trained a one dimensional CNN to classify sentences describing stress to either cellular stress or system stress (Fig. 2). To create a training corpus, we used a word2vec embeddings library based on PubMed and PubMedCentral data (Moen and Ananiadou 2013) by retrieving words that are similar to examples of systemic stress and cellular stress (e.g, restraint, corticosterone, CRH, and oxidative stress respectively). We then manually crafted two PubMed queries to retrieve abstracts related to systemic or cellular stress:

A. (CRF OR AVP OR urocortin OR vasopressin OR CRH OR restraint OR stressor OR tail-shock OR (social AND defeat) OR (foot AND shock) OR immobilization OR (predator AND odor) OR intruder OR unescapable OR inescapable OR CORT OR corticosterone OR cortisol or ACTH OR prolactin OR PRL OR adrenocorticotropin OR adrenocorticotrophin) AND stress NOT (ROS OR oxidative OR redox-regulation OR nitrosative OR nitrative OR hyperglycemia OR carbonyl OR lipoxidative OR Nrf2-driven OR thiol-oxidative)
B. (ROS OR oxidative OR redox-regulation OR nitrosative OR nitrative OR hyperglycemia OR carbonyl OR lipoxidative OR Nrf2-driven OR thiol-oxidative) AND stress

**Figure 2:**
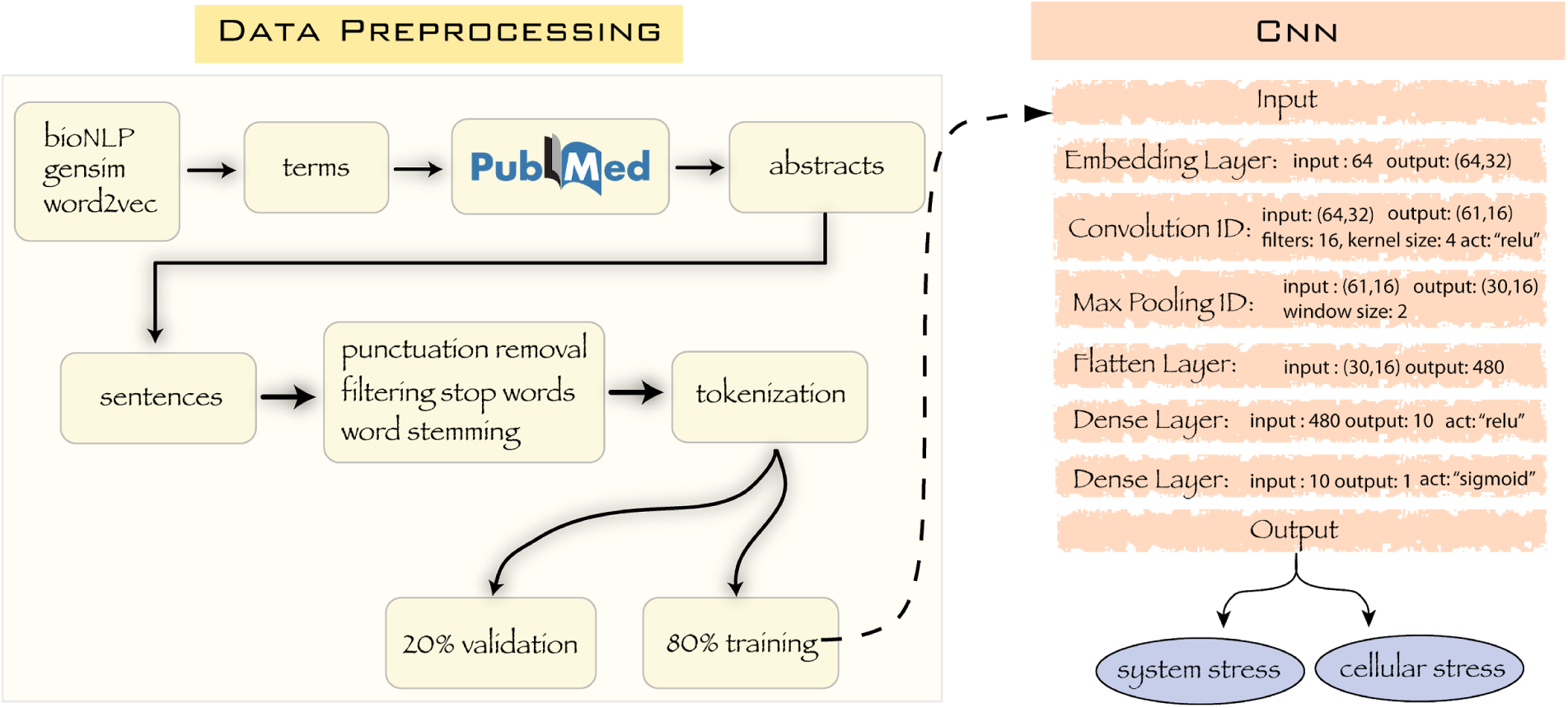
Pipeline for training the convolutional neural network that classifies sentences containing the word “stress”. Pipeline for training the convolutional neural network that classifies sentences containing the word “stress”. We used biomedical natural language processing tool (Moen and Ananiadou 2013) and the word2vec embeddings derived from PubMed and PMC text. The relevant terms with “system stress” and “cellular stress” were searched by using the cosine similarity tool in Python’s Gensim library and the abstracts including these terms were fetched from PubMed. Abstracts then were parsed into sentences, punctuations were then removed, stop words were filtered, and all words were reduced to their stems. These words were then “tokenized” and were splitted into training (80%) and validation (20%) sets. Input layer of the model passed the training data to the embedding layer, which produced a 32 dimensional embedding vector for each word. After a 1D convolutional layer with 16 filters and a kernel size of 4, downsampling is implemented by a maximum pooling layer with window size of 2. Output of this is flattened to a 480 node layer and connected to two fully connected layers. We use the rectifier unit function to activate the neurons in the convolution layer and the dense layer. Last dense layer is activated by the sigmoid function. The final weights of the model were used to classify input sentences into either system stress or cellular stress.

We downloaded all the PubMed abstracts returned from these two queries. Manually examining some of the abstracts confirmed the relevance of the results. We then extracted all sentences containing the word *stress* from each set and kept 9,974 sentences from the “systemic stress” class and 9,652 sentences from the “cellular stress” class as our stress training/validation corpus. We maintained another set of 10,000 sentences as the testing corpus, 5,000 sentences for each class.

To clean the data and make it ready for deep learning, we split 19,626 sentences into words, removed punctuation marks, filtered the stop words and stemmed the words (Brownlee 2017). The words formed a vocabulary of size 23,153 and were tokenized by the Tokenizer library of Keras API. Then the tokenized sentences were split randomly into training and validation sets at 80% and 20%, respectively. We built a 1D convolutional neural network in Keras on top of the Tensorflow framework (Abadi *et al*. 2016). The model includes an embedding layer that projects each word to a 32 dimensional space; hence this layer produces a weight matrix with 23,153 x 32 dimensions. Sentences are padded to 64 words, resulting in 64×32 sized matrices in the model. After that, a one dimensional convolutional layer with 16 filters and a kernel size of 4 is implemented and activated by the rectified linear unit (ReLU). This layer produces a 4 x 32 x 16 weight matrix. Downsampling is performed by max pooling with a window size of 2. Then a flattened layer with 480 neurons is connected to two fully connected layers, one of which has 10 neurons activated with ReLU and the latter one is the final layer activated with a sigmoid function. We validate the model using 3,924 sentences, 1,997 of them belong to the “systemic stress” class, 1,927 sentences belong to the “cellular stress” class. These were selected randomly before training. To minimize the value of the loss function and update the parameters, Adamax optimization algorithm (Kingma and Ba 2014) was used with the parameters of learning rate=0.002, beta1=0.9, beta2=0.999. The binary cross entropy loss function is used for this binary classification task. These hyperparameters were optimized using the training corpus.

We used the confusion matrix to evaluate the performance of the classification and summarize the results for the test dataset (Table 1). The rows and the columns of the matrix represent the values for the actual class and predicted class, respectively. The measures of accuracy in the table were calculated by using the values in the table; the number of true positives (TP), false negatives (FN), false positives (FP) and true negatives (TN). Sensitivity, i.e., the ratio of TP to TP+FN, is the proportion of the systemic stress sentences correctly identified. Specificity, i.e., the ratio of TN to TN+FP, is the ability of the model to identify the cellular stress sentences correctly. Precision is the proportion of the correct systemic stress sentences in the predicted class of systemic stress sentences, and is calculated as the ratio of TP to TP+FP. Accuracy of the model is the proportion of the total number predictions that are correct, and is calculated as the ratio of TP+TN to all. The performance measures including the area under the ROC curve (sensitivity vs. 1-specificity) produced by these values are given in the Results section.

**Table 1:**
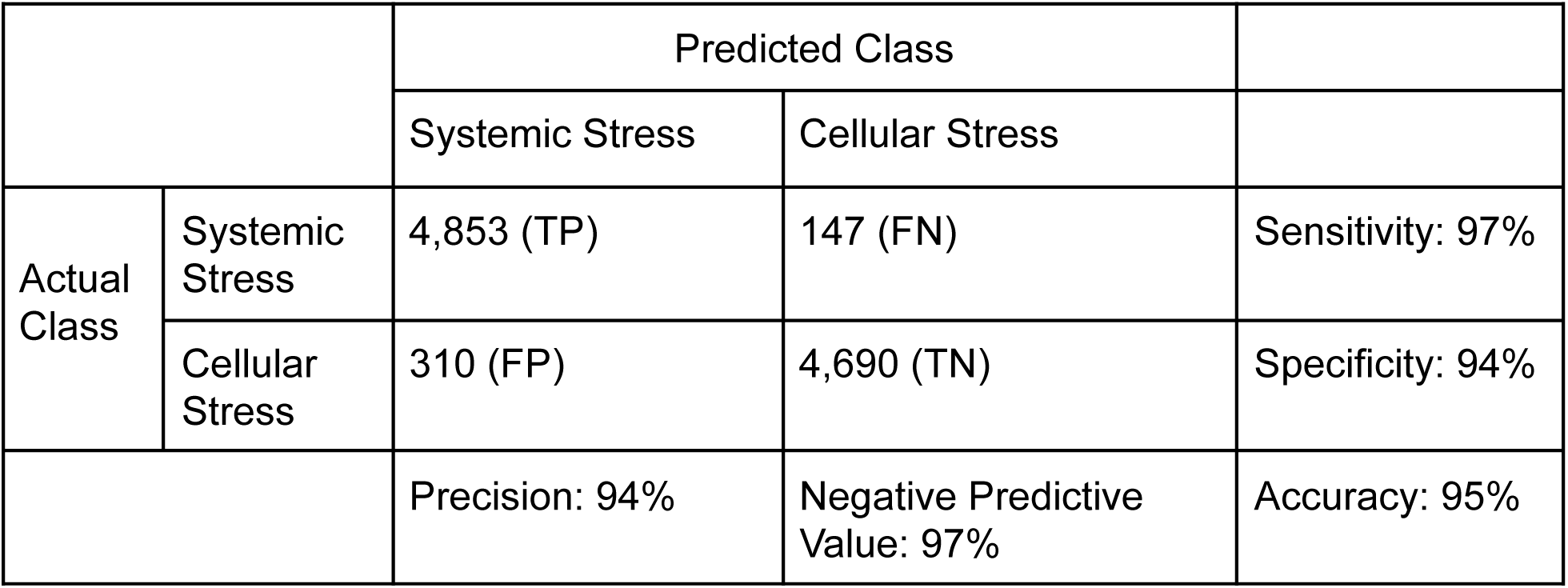
Confusion Matrix of CNN on Test Data.

## 3. Results

We have written a command line and a graphical interface for searching the role genes play in biological systems. The command line interface is more suitable for searching a large number of genes and requires the user to install the software and maintain a local mirror of PubMed. Using this interface, we queried all human genes against PubMed and identified the most researched genes in addiction. The top 10 genes with the greatest number of addiction related abstracts are FOS, BDNF, TH, OPRM1, CNR1, DRD2, CREB1, SLC6A4, TNF and CYP2B6. Many of these genes are involved in the activation of neurons or neurotransmission. In addition, some of the genes involved in the immune system function and intracellular signalling, such as TNF and IL6 are among the top genes. This list of genes are provided in Table S2. These genes and their associated sentences are available at the http://genecup.org website.

The graphical interface, on the other hand, is more user friendly and can be used through our website. A query of 3 terms can be completed in about 20–30 seconds. The query time increases linearly by the number of terms. Thus a search of 20 genes can be completed in about 2–3 minutes. Most of the time is spent on interacting with PubMed to obtain PMIDs. As a demonstration of the utility of the web interface, we entered the nine genes that reached suggestive significance in a recent genome wide association study of opioid cessation (Cox *et al*. 2020). The graph view of the search results are shown in Figure 3. Genes and keywords are all shown as circles and lines connecting them show the number of abstracts containing the two circles they connect. Keywords under the same main category are shown with the same color in the graphic output. Clicking on the lines brings up a new page that displays all sentences containing the keywords that line connect. An alternative tabular view of the same results is also available, where genes, the keywords, and number of abstracts are shown as separate columns.

**Figure 3:**
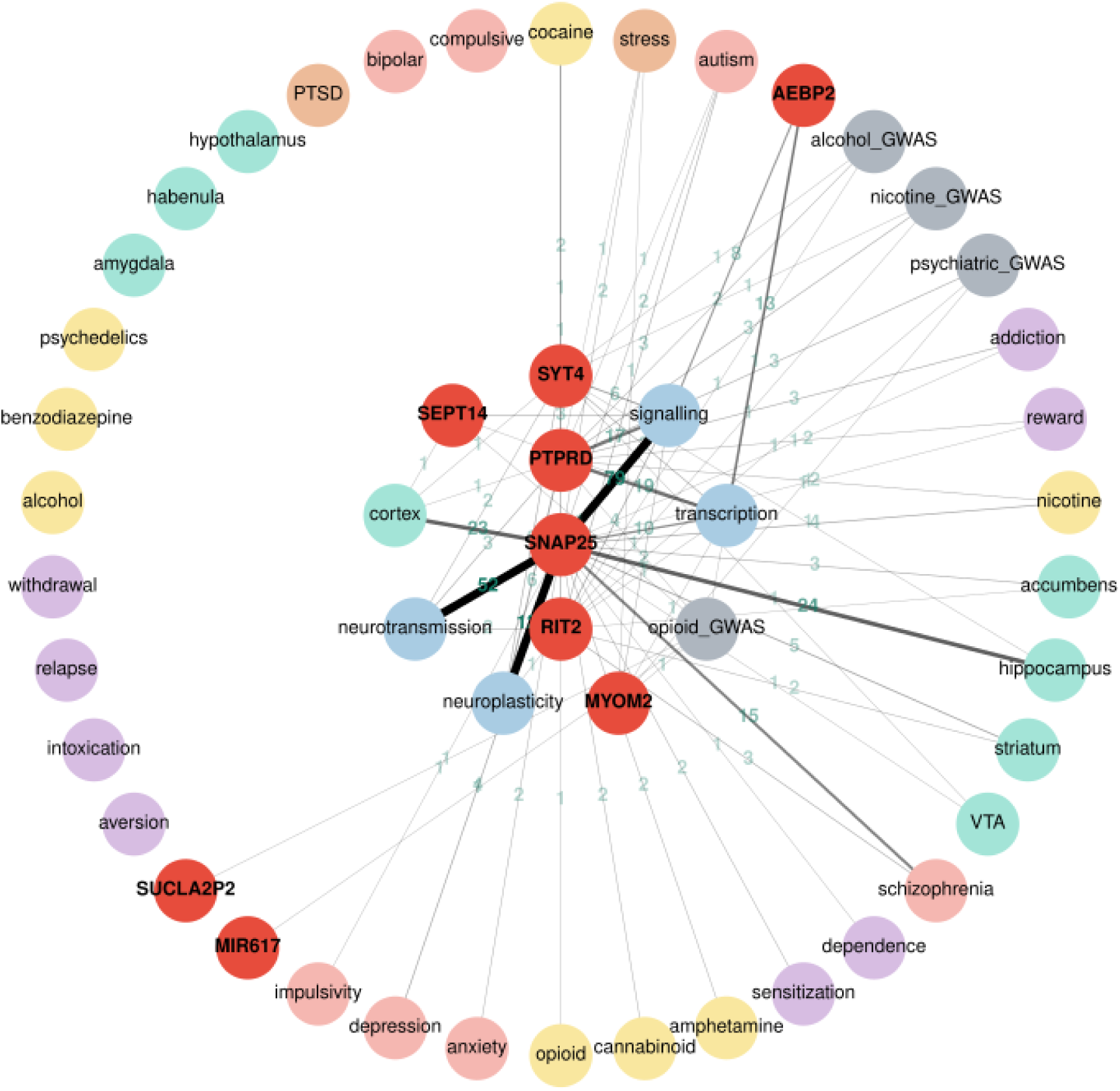
An interactive Cytoscape graph visualizing gene-keyword relationships. An interactive Cytoscape graph visualizing gene-keyword relationships. Nodes (circles) represent either search terms (in red) or keywords (colored according to the mini ontology; GWAS results are in grey). Clicking the keyword nodes displays the individual terms that are included in the search. Clicking the gene symbols displays their synonyms. The edges represent relationships between nodes. The number of PubMed abstracts where the gene symbol and keyword co-occur in the same sentence are displayed on the edges. The width of the edge is correlated with the number of abstracts. Clicking on the edges shows these sentences, which are linked back to PubMed abstracts. Nodes can be moved about for better visibility of relationships. These genes were taken from a recent genome wide association study of opioid cessation (Cox *et al*. 2020).

Our results contained sentences in PubMed that described the roles played by PTPRD, SNAP25 and MYOM2 in addiction, which were all discussed in the original publication (Cox *et al*. 2020). In addition, our results found sentences indicated the potential involvement of RIT2 and SYT4 in addiction. For example, RIT2 is associated with smoking initiation (Liu *et al*. 2019) and autism (Liu *et al*. 2016). Recent publications indicated that RIT2 is involved in dopamine transporter trafficking (Fagan *et al*. 2020) and plays a sex-specific role in acute cocaine response (Sweeney *et al*. 2020). SYT4 is expressed in the hippocampus and entorhinal cortex (Crispino *et al*. 1999) and regulates synaptic growth (Harris *et al*. 2016; Ó’Léime *et al*. 2018). Further, SUCLA2P2 has been implicated in age of smoking initiation (Argos *et al*. 2014) and Schizophrenia (Ikeda *et al*. 2019). This example demonstrated the utility of GeneCup in rapidly finding information that links a gene to addiction and thus integrating new findings with previous research findings.

For sentences containing the word “stress”, we designed a one-dimensional convolutional neural network with 4 hidden layers (Fig. 2) to differentiate them into two classes, namely systemic and cellular stress. The neural network was optimized using the gradient based optimization algorithm Adamax. During training, model accuracy (Fig. S1.A) increased rapidly during the first five epochs to approximately 0.995, while validation accuracy peaked at 0.991 at epoch five. On the other hand, model loss curve (Fig. S1.B) on the training dataset continued to decline after the initial drop and approached zero after 15 epochs. However, the loss on the validation data set started to increase after epoch five, indication model overfitting. Therefore, we used the weights that maximized the validation performance before overfitting (i.e. epoch five). By using these weights and parameters, our model has an AUC of 99.2% on the validation dataset.

We tested the model on a new dataset consisting of 5,000 system stress sentences and 5,000 cellular stress sentences. The confusion matrix for the prediction of the test dataset is presented as Table 1. The sensitivity of the model that is the proportion of predicted systemic class sentences to all sentences observed in this class is 97%. The similar measure for cellular class sentences, i.e., specificity is 94%. The prediction accuracy of the model that is the ability to distinguish two classes on the test dataset is 95.4% and the AUC is 98.9% for the test dataset.

We also checked the distribution of the predicted probabilities (Fig. S2) of the test dataset. The model predicts a probability of the class membership for each sentence. If the predicted probability of a sentence is more than 0.5, it is labelled as a system stress sentence. Otherwise the sentence is predicted to be a member of the cellular stress class. Among the system stress sentences in the test dataset, 88% of the sentences had predicted probabilities greater than 0.9. This shows the model’s confidence of its prediction on stress sentences. Likewise, 88% of the cellular stress sentences had predicted probabilities less than 0.1. Therefore the model is 90% confident about the classification of 88% of the cellular stress sentences.

The weights of the trained model are saved on the server and are used to make predictions for each retrieved sentence when the user clicks on the edge connecting the stress category and the gene name (Fig. 4). As an example of run time performance, it took approximately 12 seconds to classify 3,908 sentences on CRF and stress.

**Figure 4.**
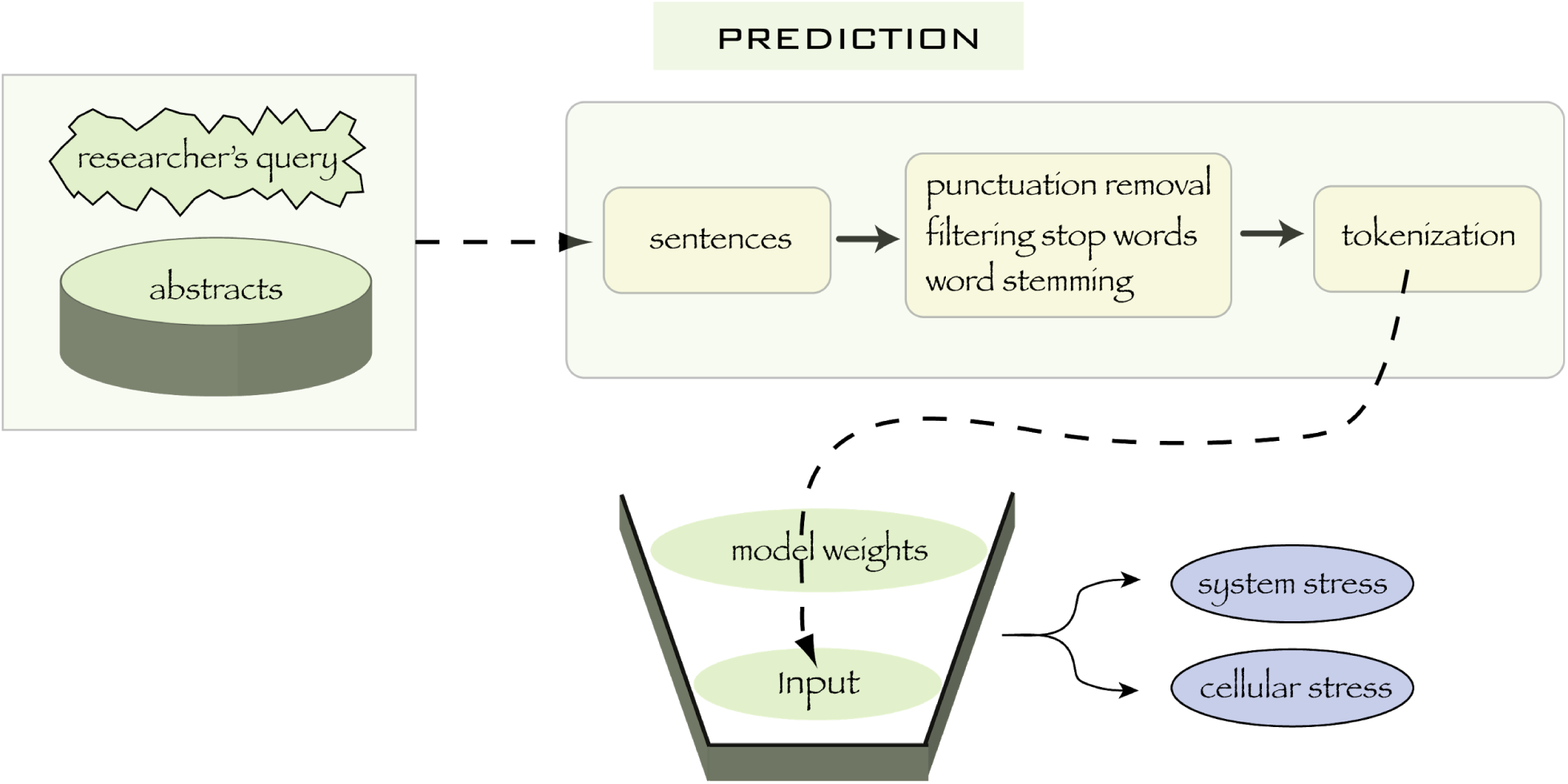
Steps for classifying sentences using a trained neural network. Steps for classifying sentences using a trained neural network. Abstracts are fetched from the locally mirrored copy of PubMed and are parsed into sentences. Punctuation marks and stop words are removed and the remaining words of the sentences are stemmed. The words are tokenized by using the Tokenizer library of the Keras API. The weight matrices of the trained model are multiplied by the sentence matrix to predict whether the input sentences are related to system stress or cellular stress.

## 4. Discussion

We present here a literature mining web application, GeneCup, that extracts sentences from a locally mirrored copy of PubMed abstracts containing user provided gene symbols and the keywords of the custom ontology. Associations between the genes and various phenotypes from human GWAS results are also provided. The users can include up to 200 gene symbols in each search. The results are presented in a graphical or a tabular format, both provide links to review individual sentences that contain the gene and at least one keyword. Gene synonyms are also presented and can be included in additional searches. We also provide an addiction ontology that is approximately 300 predefined addiction-related keywords organized into seven categories. Stress related sentences are automatically classified into system vs cellular stress if the addiction ontology is used.

Scientists using omics methods face a particularly challenging task when trying to integrate new findings with existing knowledge. The increasing number of genes contained in data sets, the breadth of sciences, and the large amount of existing knowledge captured in PubMed make systematic literature surveys daunting tasks. Typically, scientists manually conduct more detailed searches in areas where they have expertise and the queries are much less thorough in other areas. The search strategies are often crafted ad hoc and likely different from one day to another.

GeneCup provides an interface that allows comprehensive queries of the role of any gene using a set of user defined keywords. Although most of the functions provided by GeneCup can be carried out manually, it will require several orders of magnitude more time and effort. Even then, the manually collected results will be difficult to review. In contrast, results provided by GeneCup are automatically organized by the ontology. All the genes and keywords can be seen in one graph or table, with informative sentences and abstracts readily available.

We also curated an addiction ontology of about 300 keywords. These keywords provide a comprehensive coverage of key concepts related to addiction. The applied machine learning solution to resolve the ambiguity of the word stress further reduced the burden of the user when coming through the vast amount of literature on stress.

GeneCup presents to the user sentences containing genes and keywords of interest to the user. Compared to phrases or abstracts, sentences are the most succinct semantic unit to convey a fact. Ding et al (2002) compared different text processing units for text mining system design and found that the highest precision of information retrieval is achieved when phrases are used as the text unit whereas using sentences are more effective than both phrases and abstracts. Therefore, similar to our previous text mining tool (Chen and Sharp 2004), we continue to use sentences as the information unit. Unlike the commonly used gene ontology enrichment (Osborne *et al*. 2007) or gene set enrichment (Subramanian *et al*. 2005) analysis, the literature analysis provided by GeneCup does not evaluate any statistical significance. Instead, these key sentences provide easy access to relevant prior research, where the nuanced details can be easily obtained by following the link from the sentence to the abstract and then to the full text article.

Stress plays key roles in addiction. Using a convolutional network, we trained a model that achieved 97% sensitivity and 94% specificity in classifying sentences containing the word stress to either systemic stress or cellular stress. Training such a model requires large amounts of labeled data. Manually labeling these data is very labor intensive. Using an approach that is similar to some recent advances in automated data labeling (Ratner *et al*. 2020), we carefully crafted two PubMed queries to obtain over 30,000 sentences that mostly belong to the correct category. This large corpus of text allowed us to achieve peak classification performance with less than 5 epochs of training (Fig. S1).

Gene synonyms represent a large challenge to any text mining approach. Not including synonyms will result in the loss of information. However, many synonyms, especially those that are short, have multiple meanings. For example, CNR is a synonym for the CNR1 gene. However, CNR is also an acronym for contrast noise ratio, frequently used in imaging analysis literature. We manually edited the list of aliases for the most studied 100 addiction related genes, which are shown in Table S2. For user supplied gene symbols, we do not include synonyms in the initial search to prevent the noise from “drowning out” the signal. However, we do provide users an option to either search individual synonyms or to conduct a combined search of all synonyms as a secondary step. We think this middle-of-the-road approach is the most efficient method to achieve a balance between computation and performance. Future work can potentially use deep learning to classify all PubMed abstracts for their relevance to addiction and thus exclude many abstracts containing short words that are not relevant to addiction from being confounded with gene synonyms.

Other future improvements for GeneCup are possible. For example, GeneCup uses PubMed abstracts as the source of data, rather than PubMed Central, which contains full-text articles. Lin (Lin 2009) compared the effectiveness of information retrieval from abstract vs full text search and found that full text search, when indexed using paragraphs as the unit, is more effective than the abstract-only search. Several groups have reported either using full text search for curation (Van Auken *et al*. 2014; Müller *et al*. 2018) or using full text for analysis (Wei and Collier 2011; Verspoor *et al*. 2012; Islamaj Dogan *et al*. 2017). NCBI also provides an API for PubMed Central. However, the majority of the articles in PubMed Central are subject to traditional copyright restriction (“PMC Open Access” 2020) and it is not feasible to establish a local mirror of the full-text collection. Interactively retrieving text via NCBI API is not feasible on the scale we need (e.g, several thousand articles at a time). Further, we anticipate full text may cause duplications of information and increase the noise in results.

GeneCup does not retrieve relationships between genes. There are several existing tools available for this purpose, such as Chilibot (Chen and Sharp 2004), or GeneMania (Warde-Farley *et al*. 2010). Instead, GeneCup focuses on the relationship between genes and a set of keywords organized as an ontology. The addiction ontology was developed based on the expertise of the authors. It certainly contains biases and can be further improved. For example, tight integration with community developed ontology for addiction or psychiatric disease, such as those that are available from the Open Biological and Biomedical Ontology Foundry (www.obofoundry.org), or automated methods for converting MESH headings can be tested in the future.

## Availability of data and materials

GeneCup is a free and open source web application. The source code of GeneCup and the link to a running instance is available at https://github.com/hakangunturkun/GeneCup.

## Authors contribution

MHG conducted the research and drafted the manuscript. HC conceived of the project and conducted the initial research. EF, TW, MKM, RWW and PP contributed to the research. All authors revised and approved the final manuscript.

---

**Supplementary Figure 1:**
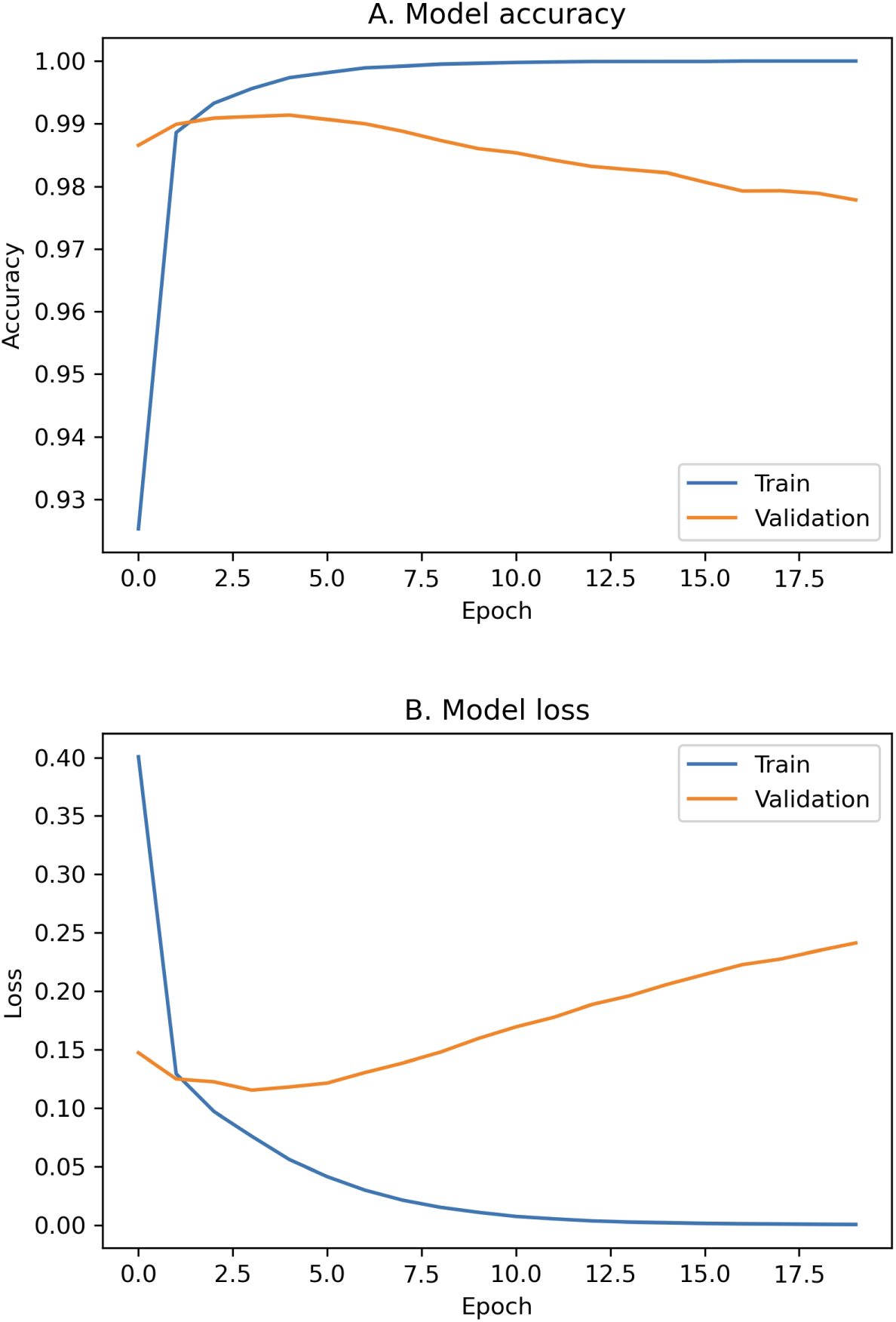
The accuracy and loss curve of a convolutional neural network trains to classify sentences containing the word “stress”. A convolutional neural network was trained on a dataset containing 19,626 sentences. This dataset was splitted into two parts, having 80% for training and 20% for validation. The gradient based Adamax algorithm was deployed with a learning rate of 0.002 during model training. The accuracy (A) increased rapidly up to 0.995 after the first five epochs. At the same time, the validation accuracy was maximized at 0.991. On the other hand, the loss curve (B) experienced a sharp fall followed by a continuous decrease. The increase of the loss curve on the validation set after the fifth epoch was an indication of an overfitting. To avoid overfitting we used the parameters that maximized the validation performance. Our model with these parameters has an AUC of 99.2%.

**Supplementary Figure 2:**
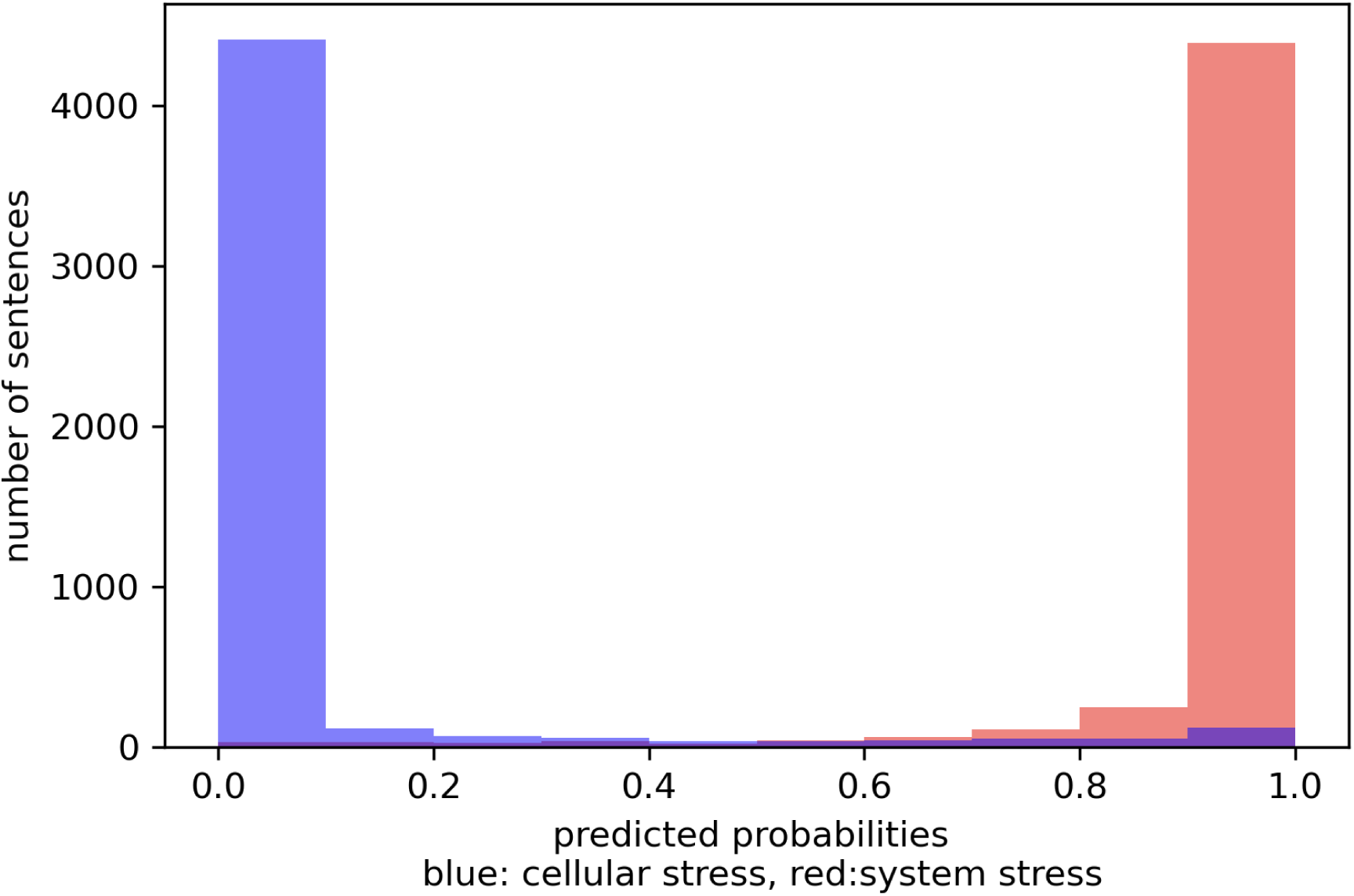
Distribution of the predicted probabilities of the test dataset. We tested the convolutional neural network model on a dataset including 5000 sentences from each class containing the cellular stress and system stress related sentences. In order to have a better understanding on the model’s reliability of its prediction on the new data, we checked the distribution of the predicted probabilities of the test dataset. The bars represent the number of sentences having the predicted probabilities shown on the x-axis (histogram). The sentences having predicted probabilities greater than 0.5 are labelled as systemic stress sentences (red bars). The blue bars represent sentences belonging to cellular stress class. Among the system stress sentences in the test dataset, 88% of them had predicted probabilities greater than 0.9. Similarly 88% of the cellular stress sentences had predicted probabilities less than 0.1. This indicates that the model has a 90% confidence about the classification of the 88% of the cellular stress sentences.

**Supplementary Figure 3:**
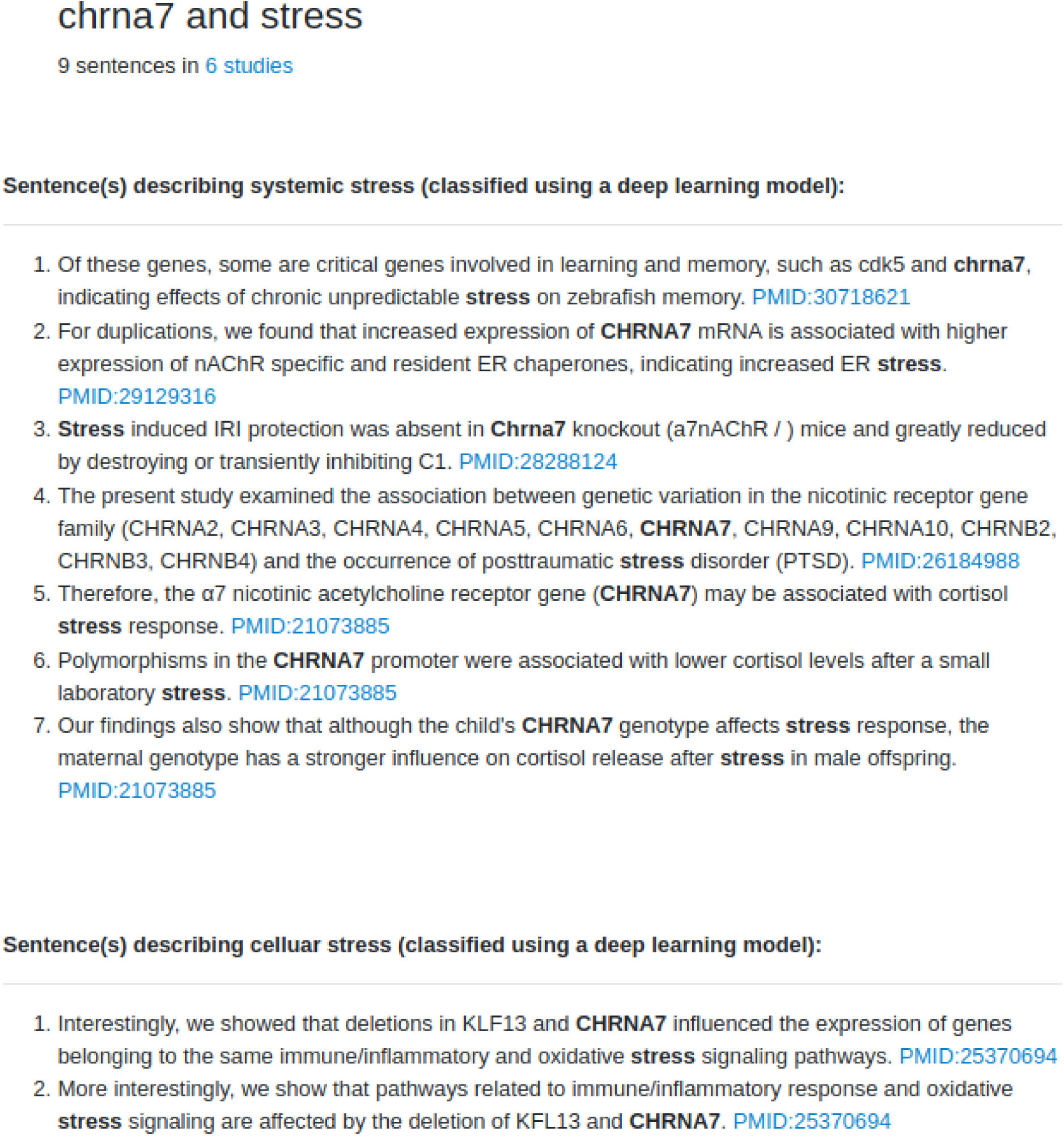
One output page in GeneCup displaying sentences retrieved when querying *chrna7* against the keyword *stress*. The sentences describing stress are classified according to its relation to systemic stress or cellular stress by a convolutional neural network. GeneCup retrieved 9 sentences containing chrna7 and stress from 6 studies. Among them, 7 sentences in 5 studies are predicted as a systemic stress sentence and 2 sentences in 1 study are predicted as cellular stress sentences.

**Supplementary Table 1.**
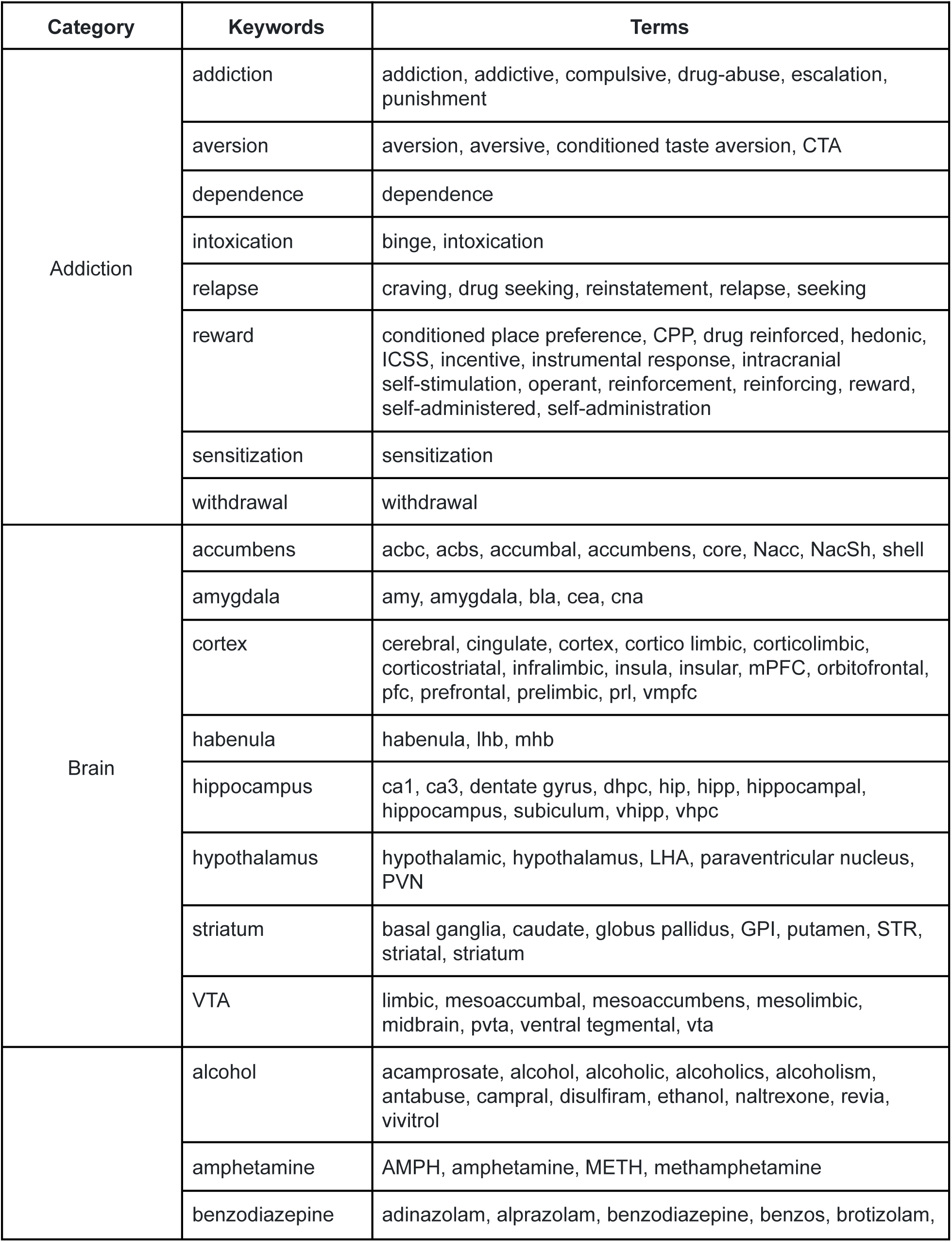
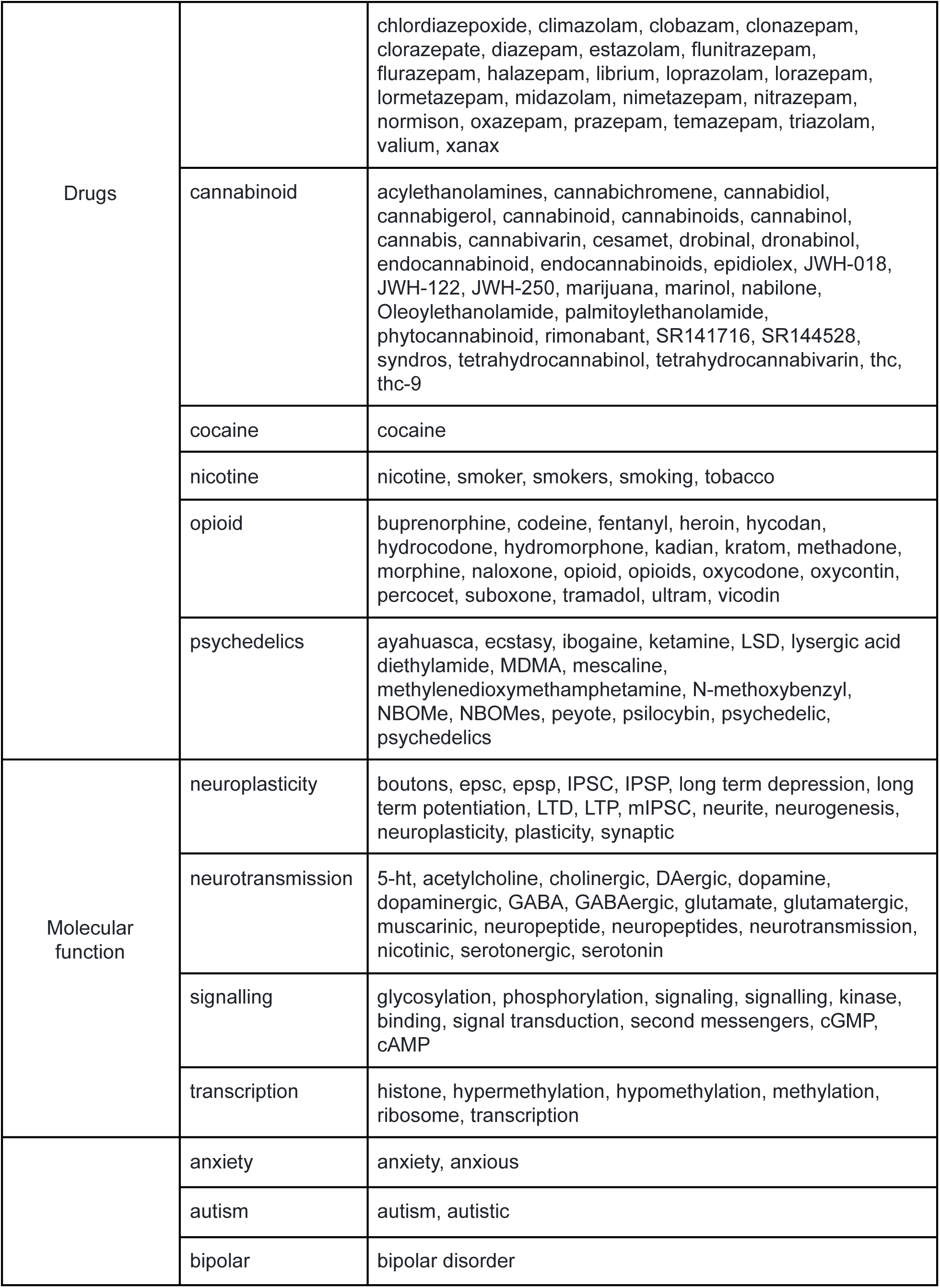
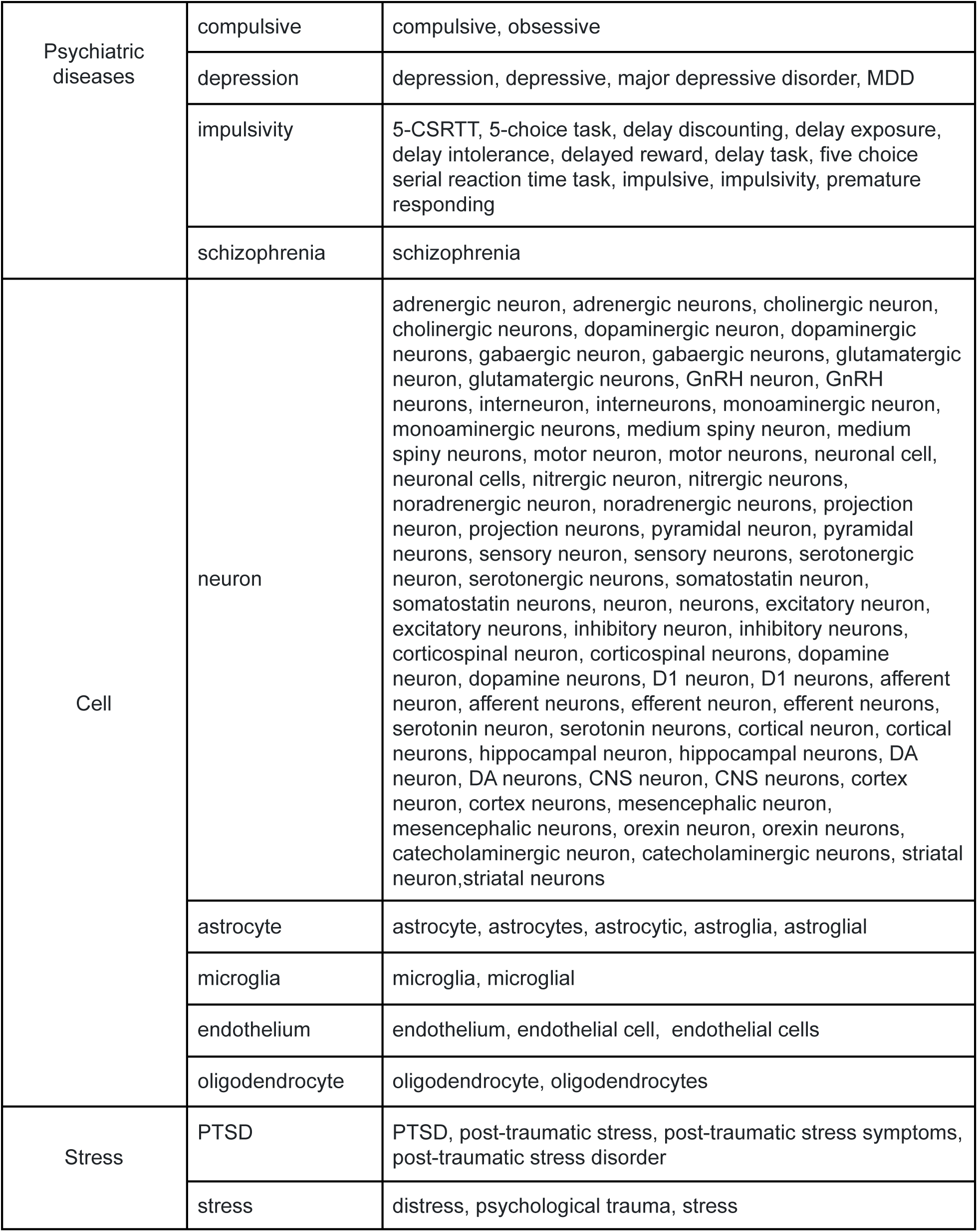
Mini ontology for addiction related concepts

**Supplementary Table 2.**
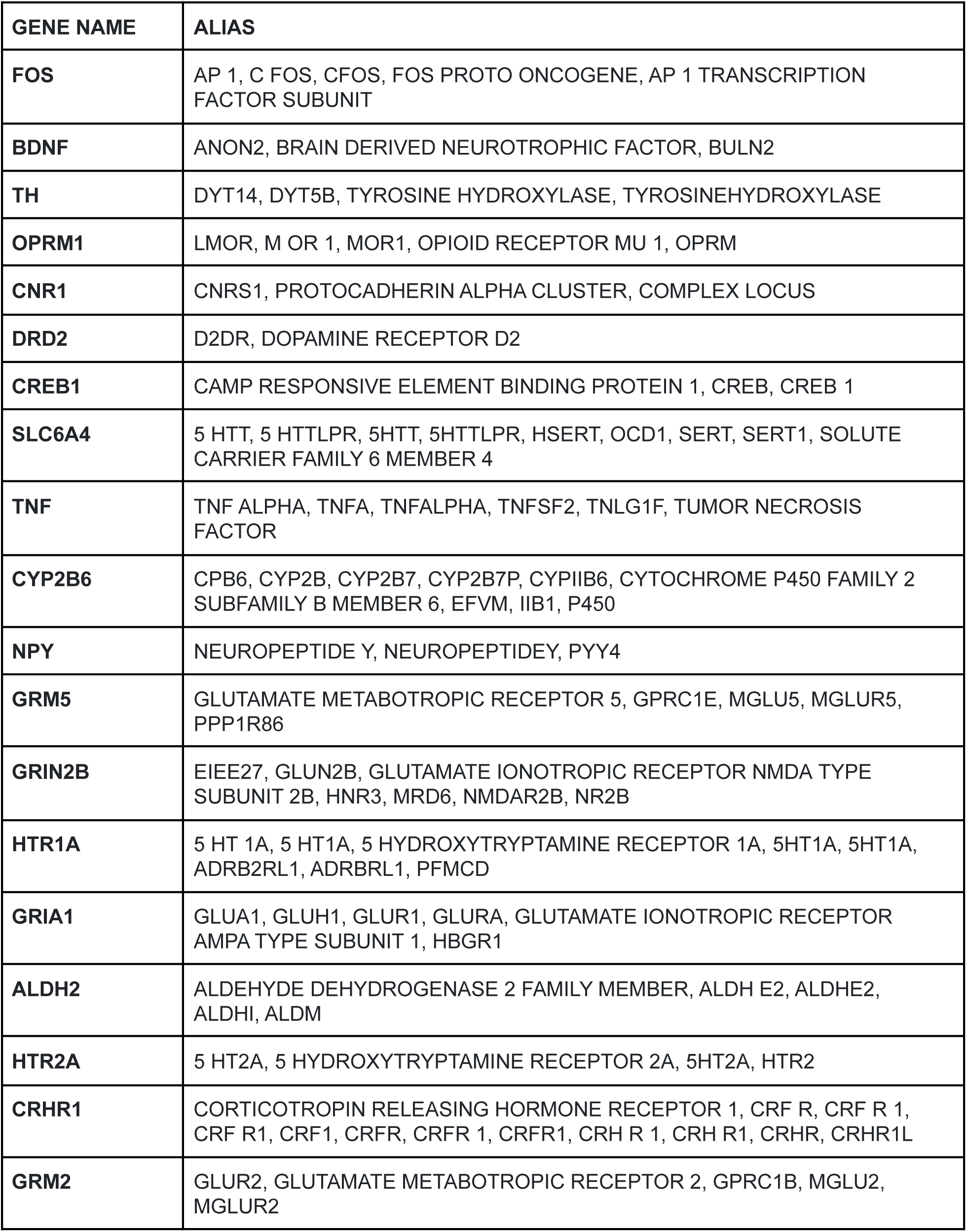
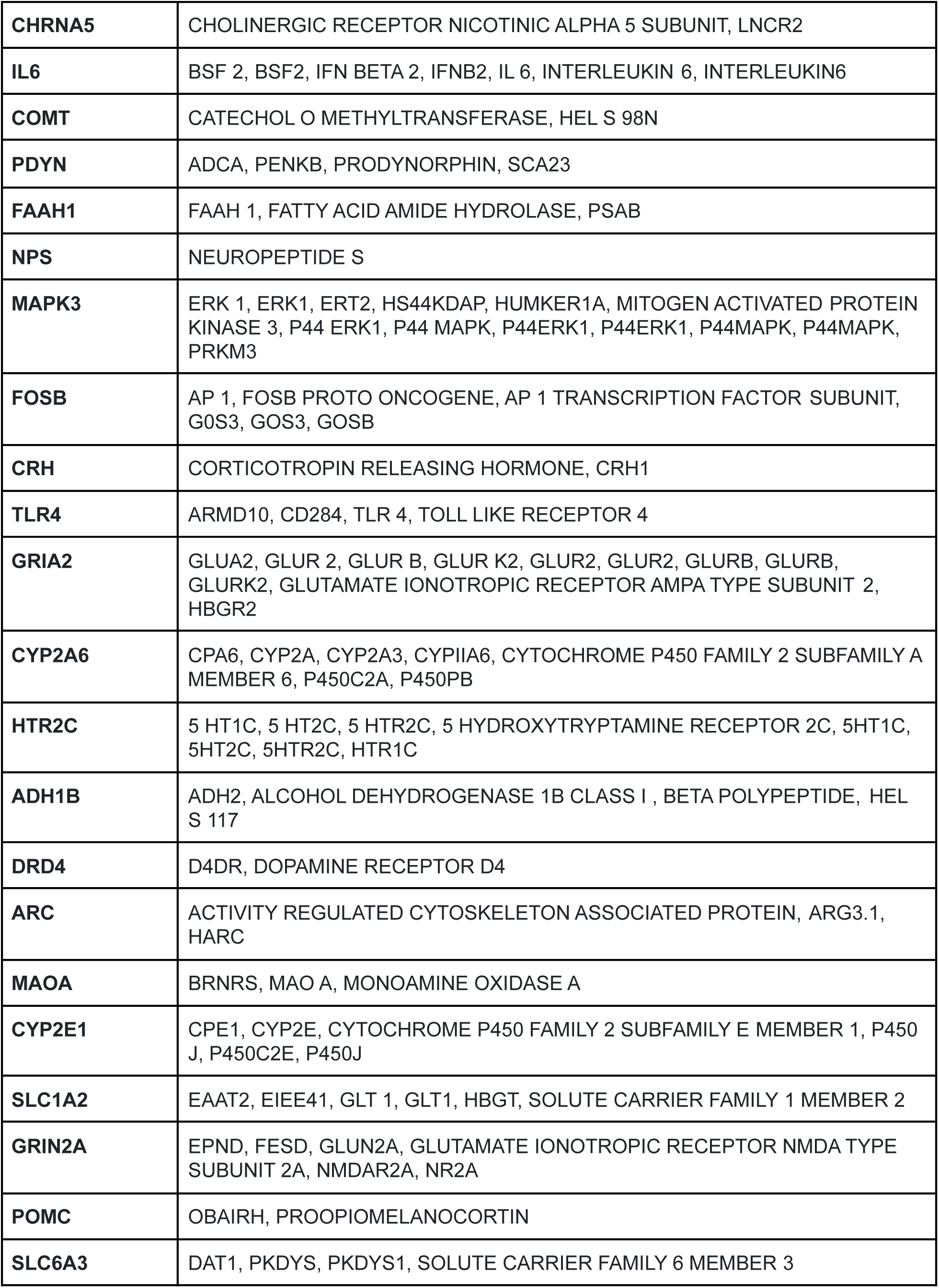
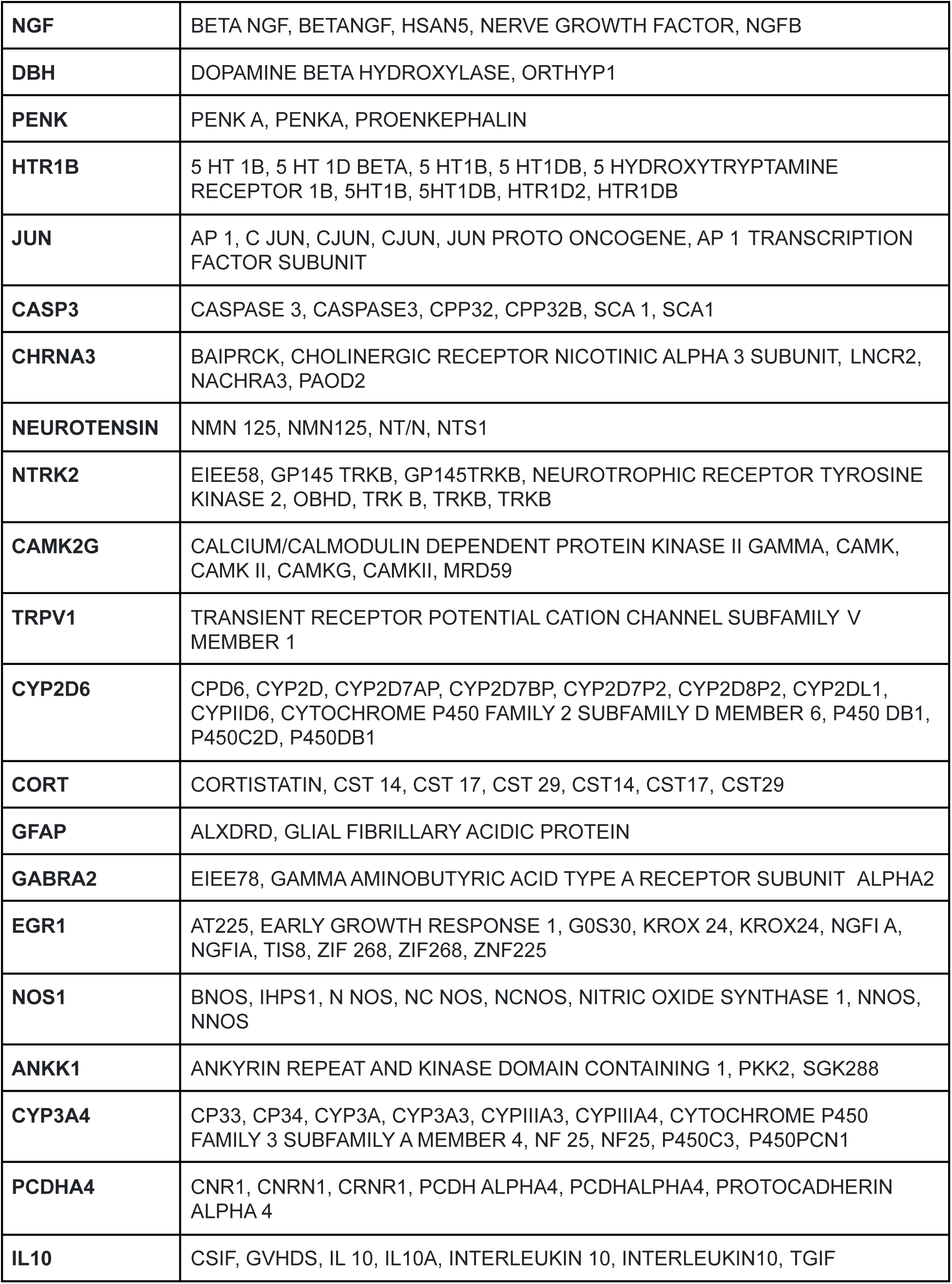
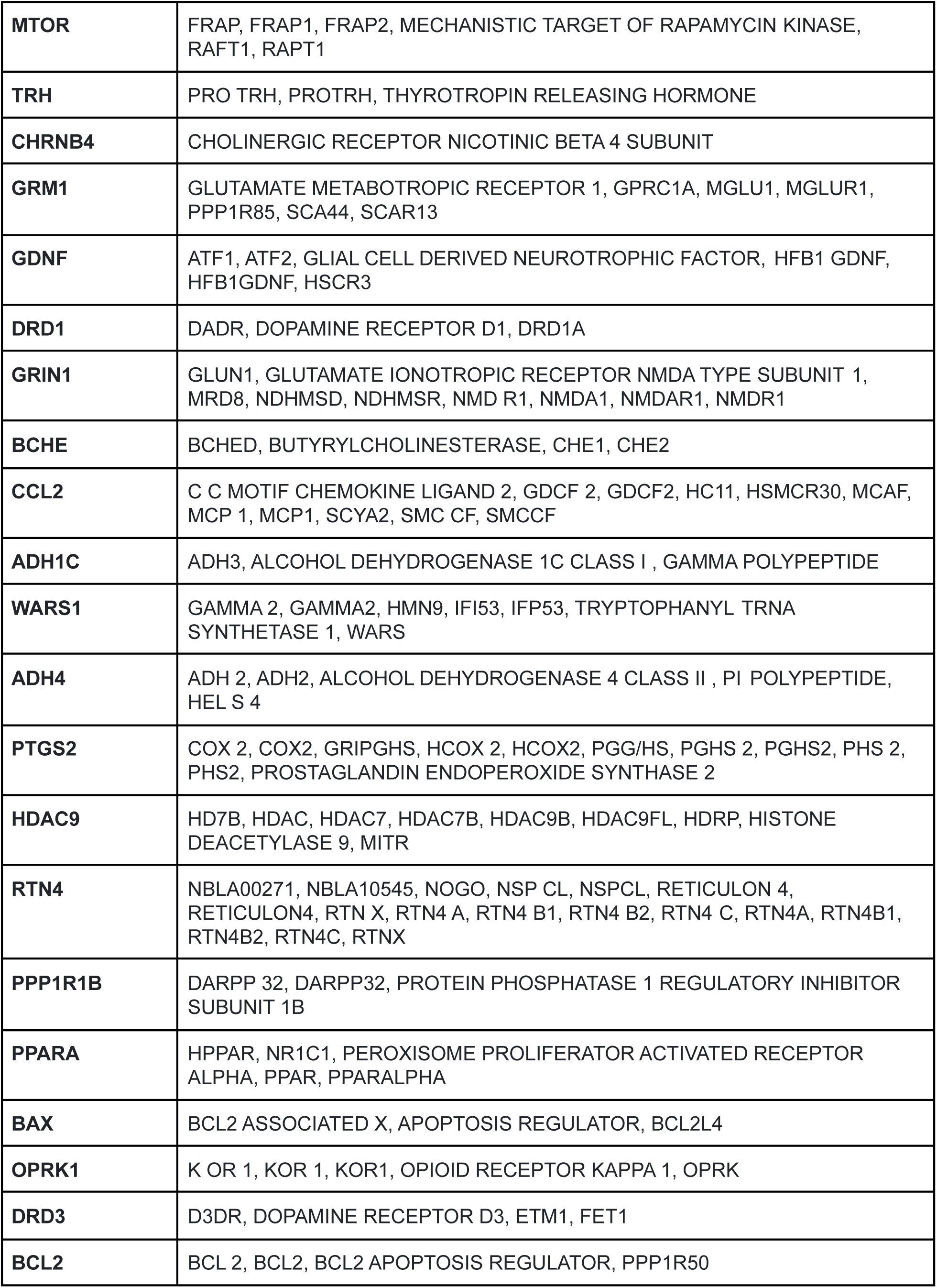
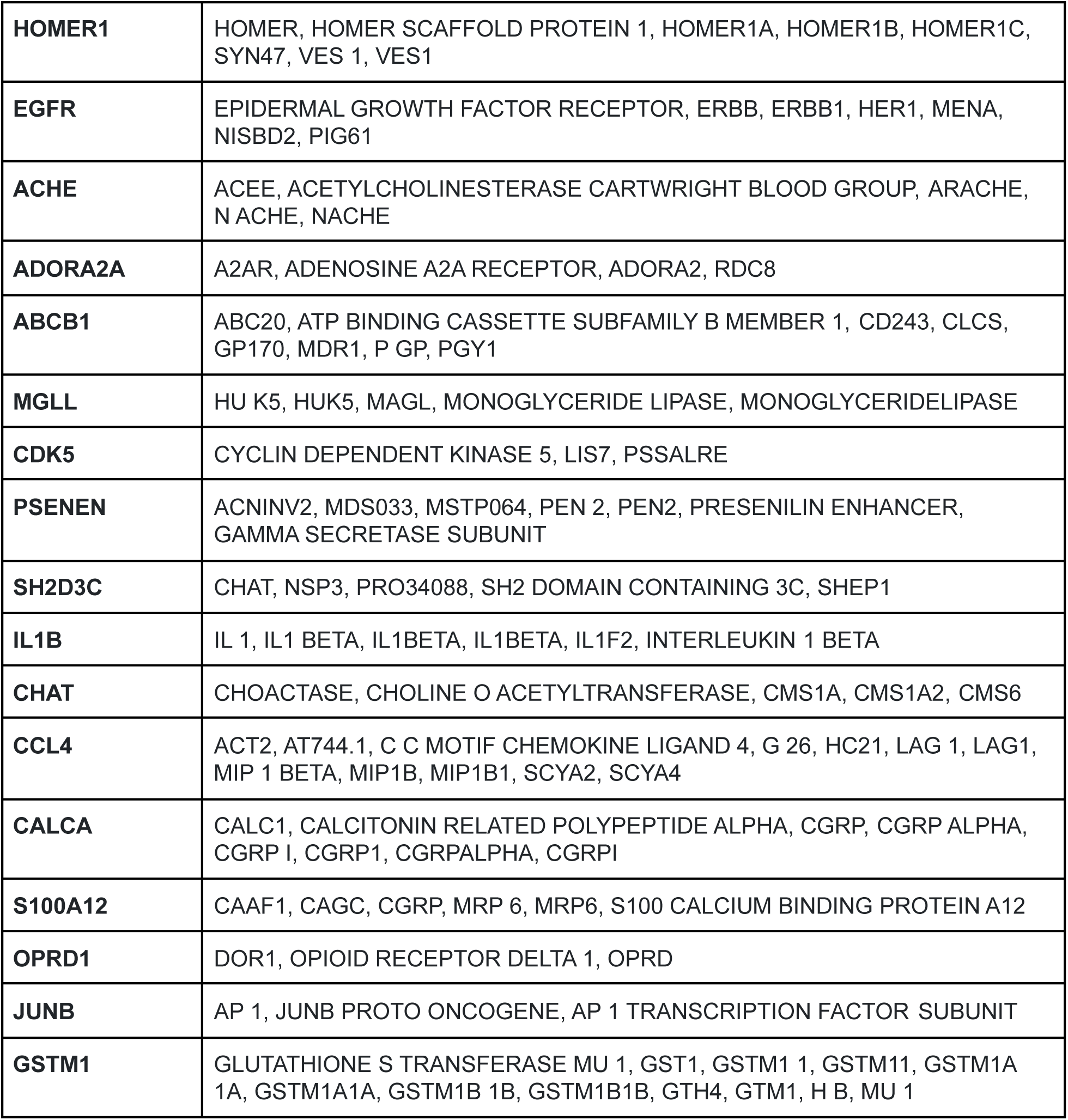
Most studied addiction related genes and their aliases

